# *De novo* transcriptome sequencing of the northern fowl mite, *Ornithonyssus sylviarum*, shed light into parasitiform poultry mites evolution and its chemoreceptor repertoires

**DOI:** 10.1101/2021.03.17.435780

**Authors:** Biswajit Bhowmick, Huaqing Chen, Jesus Lozano-Fernandez, Joel Vizueta, Rickard Ignell, Qian Han

## Abstract

The poultry red mite (PRM), *Dermanyssus gallinae* (De Geer), and the northern fowl mite (NFM), *Ornithonyssus sylviarum* (Canestrini and Fanzago), are the most serious pests of poultry, both of which have an expanding global prevalence. Research on NFM has been constrained by a lack of genomic and transcriptomic data. Here, we report and analyze the first transcriptome data for this species. A total of 28,999 unigenes were assembled, of which 19,750 (68.10%) were annotated using seven functional databases. The biological function of these unigenes was predicted using the GO, KOG, and KEGG databases. To gain insight into the chemosensory receptor-based system of parasitiform mites, we furthermore assessed the gene repertoire of gustatory receptors (GRs) and ionotropic receptors (IRs), both of which encode putative ligand-gated ion channel proteins. While these receptors are well characterized in insect model species, our understanding of chemosensory detection in mites and ticks is in its infancy. To address this paucity of data, we identified 9 IR/iGluRs and 2 GRs genes by analyzing transcriptome data in the NFM, while 9 GRs and 41 IR/iGluRs genes were annotated in the PRM genome. Taken together, the transcriptomic and genomic annotation of these two species provide a valuable reference for studies of parasitiform mites, and also helps to understand how chemosensory gene family expansion/contraction events may have been reshaped by an obligate parasitic lifestyle compared with their free-living closest relatives. Future studies should include additional species to validate this observation, and functional characterization of the identified proteins as a step forward in identifying tools for controlling these poultry pests.

## Introduction

The northern fowl mite (NFM), *Ornithonyssus sylviarum*, and the poultry red mite (PRM), *Dermanyssus gallinae* (Parasitiformes: Mesostigmata: Dermanyssoidea), are small (< 1 mm) obligate blood-feeding ectoparasites of birds that cause significant economic losses to the poultry industry, and a negative impact on animal welfare. The entire life-cycle of NFM occurs on the host, while PRM lives off-host in the cracks and crevices in poultry houses (Tomley et al., 2018; Sparagano and Ho, 2020; Murillo et al., 2020). The development of novel intervention strategies to control poultry mite population is of great interest. In this context, genomic information is essential, but not available for the NFM. Until now, only a few examples of nucleotide sequences referring to the mitochondrial genome of NFM are deposited in the NCBI database (Roy et al., 2009; Bhowmick et al., 2019). With the development of high throughput sequencing (HTS) technologies, scientists have been able to study the transcriptomes of Acari (Parasitiformes and Acariformes orders), with a few reference genomes currently available. The first transcriptome analysis was carried out on PRM (Schicht et al., 2014), for which the draft genome assembly was reported based on combined PacBio and Oxford Nanopore MinION long-read *de novo* sequencing (Burgess et al., 2018).

Similar to most insects (Joseph and Carlson, 2015), non-insect arthropods, such as parasitiform ectoparasites (*Ixodes scapularis*, *O. sylviarum*, *D*. *gallinae*, and *Varroa destructor*), rely heavily on their sense of smell to discriminate and locate resources, including their vertebrate hosts (Sonenshine, 2004; Gay et al., 2020; Light et al., 2020; Faraone et al., 2020; Josek et al., 2021). To do so, non-insect arthropods seem to rely on chemosensory receptors from two gene families: gustatory receptors (GRs) and ionotropic receptors (IRs) (Josek et al., 2018). In insects, the GR family of seven-transmembrane (TM) proteins comprises up to hundreds of highly divergent (only 8–12% sequence identity) receptors, that are involved in the detection of various tastants, including e.g., bitters, sugars and cuticular pheromone compounds, as well as carbon dioxide (Vosshall et al., 1999; Clyne et al., 2000; Chyb et al., 2003; Jones et al., 2007). The IRs are members of the second largest family of chemoreceptors in insects, and represents a highly divergent subfamily of ionotropic glutamate receptors (iGluRs), which include the NMDA-type receptors, AMPA-type receptors, and kainate-type receptors that have ultra-conserved roles in synaptic transmission in animals, where the latter two are grouped together as non-NMDA types (Benton et al., 2009; Sobolevsky et al., 2009). The IRs involved in the detection of chemical stimulus (e.g. odorants), require a co-receptor (IR8a, IR25a or IR76b) to function (Abuin et al., 2011). These co-receptors along with IRs are involved in different physiological functions, such as the perception of temperature or humidity and are conserved across insects and other arthropods (Enjin et al., 2016; Ni et al., 2016).

In the present study, we performed a high-coverage sequencing and *de novo* assembly of the parasitiform mite *O*. *sylviarum*. Based on this assembly, we discuss functional categories and annotations, functional genes associated with various physiological functions, and transcriptional factors of relevant genes and gene families. Gene ontology (GO) and KEGG enrichment analyses were conducted to identify the biological functions and predict the pathways of key genes. Our whole-transcriptome data provides important molecular information on mite biology and might provide new opportunities for the development of novel control approaches against mite infestation, such as the identification of new drug targets or the development of potential vaccine candidates. The generation of the HTS data, furthermore, allowed for the investigation of chemosensory gene family evolution, such as IRs and GRs across arthropods (Eyun et al., 2017; Vizueta et al., 2020a). In both mite species, we identified and classified the putative IR/iGluR and GR unigenes using HTS data. These genes were analyzed using a series of bioinformatics tools to exhaustively understand their characteristics, including their protein structure, molecular domain architecture, and phylogenetic relationships with IR/iGluRs of other mite and tick species. In addition, we performed an automated refining and validation of the candidate gene model via ORCAE (Online Resource for Community Annotation of Eukaryotes), which offers a community-based genome annotation platform in which multiple researchers from around the world can contribute new information on gene structure, function, and gene expression atlas (Sterck et al., 2012; Yssel et al., 2019; Burgess et al., 2019). This study yields complementary information on chemosensory receptor gene families in poultry mites and other mite and tick species and provides an enriched set of gene data of traditional iGluRs, conserved IRs, and GRs, which are essential for further research on gene evolution in chemoreceptor families.

## Materials and methods

### Mite collection, RNA extraction and quality determination

The NFMs were collected from a commercial layer poultry farm, as previously described (Wenchang, Hainan, China; 19° 29’ 1” North, 110° 46’ 18” East) (Bhowmick et al., 2019). All life-stages and sexes of *O. sylviarum* mites were prepared for total RNA extraction. The samples were washed in ice-cold phosphate buffer saline (PBS) in order to clean the surfaces of the mites and then stored at −80◻ for later use. A pooled sample with multiple individuals was homogenized by snap freezing in liquid nitrogen, and then RNA extraction was performed according to the TRIzol reagent protocol (Invitrogen, CA, USA). RNA parameters and quality were assessed using an Agilent BioAnalyzer, and the concentration and purity were assessed using the NanoDrop 2000.

### cDNA library preparation and next generation sequencing

The pooled RNA sample was sent to BGI (Beijing Genomics Institute, China) for library generation and next generation sequencing (NGS). Briefly, the mRNA was isolated using the oligo (dT)-attached magnetic beads, and then a fragmentation buffer was used to break the mRNA into short fragments as templates for first-strand cDNA synthesis. The resulting first-stranded cDNA was generated using reverse transcriptase with random hexamer primer mix, and then the resulting second-stranded cDNA was synthesized using DNA polymerase I, buffer, dNTPs and RNase H. Subsequently, the synthesized double-stranded cDNA was purified and blunt end repair was carried out, adding a base A and a linker to the 3′ end. After the end repair and ligation of adaptors, the product was amplified by PCR and purified using a PCR purification kit to create a single-stranded circular DNA library. Finally, a sequencing library was performed using the BGISEQ-500 platforms.

### *De novo* transcriptome assembly, annotation, and functional classification

Prior to assembly and mapping, clean reads were obtained by removing those with low quality, adapter contamination, and poly-N from the raw reads. Meanwhile, phred score, Q20 (%), Q30 (%) and GC content in the clean reads were calculated (Cock et al., 2010). Trinity (v2.0.6) package was used to assemble the high-quality clean data, and the assembled unigenes were then analyzed using the following databases: Nr (ftp://ftp.ncbi.nlm.nih.gov/blast/db; non-redundant protein sequences), Nt (ftp://ftp.ncbi.nlm.nih.gov/blast/db; non-redundant nucleotide sequences), Pfam (http://pfam.xfam.org; protein family), KOG / COG (https://www.ncbi.nlm.nih.gov/COG/; clusters of orthologous groups), Swiss-Prot (a manually annotated and reviewed protein sequence database), KEGG (http://www.genome.jp/kegg; Kyoto Encyclopedia of Genes and Genomes), GO (http://geneontology.org; Gene Ontology), and CD-HIT (https://gvolante.riken.jp/analysis.html). In addition, TransDecoder (https://transdecoder.github.io) software was then used to determine coding regions (CDS) in unigenes. To identify genes encoding transcription factor (TF) families, all unigenes were analyzed using the Animal Transcription Factor Database (AnimalTFDB), and protein domains were identified using HMMER (Finn et al., 2015). Finally, to assess the completeness of the transcriptomes, we evaluated its content by benchmarking it against the arthropoda single-copy orthologous database using BUSCO V3 (Simão et al., 2015).

### Identification, annotation and phylogenetic analyses of IR/iGluRs and GRs

The IR/iGluR and GR gene family members were annotated in the genome of the PRM (BioProject accession number PRJNA487003; Burgess et al., 2018). Briefly, the BITACORA pipeline (Vizueta et al., 2020b) was used to conduct homology-based searches and annotate the IR/iGluR and GR genes encoded in the input genome, using a curated database of these genes enriched in chelicerate species. The output of BITACORA included gene-finding format (GFF), with both curated and newly identified gene models, and a FASTA file of predicted protein sequences (Vizueta et al., 2018). To identify the surveyed receptors in the NFM transcriptome, we then used a three-step approach to annotate the IR/iGluRs and GRs in the assembled transcripts. Firstly, IR/iGluR and GR unigenes from the transcriptomes were retrieved by the functional annotation results based on seven functional databases (NR, KEGG, KOG, NT, SwissProt, Pfam, and GO). Then, iterative tBLASTn and PSI-BLAST searches were performed with previously identified amino acid sequences of the parasitiform tick *I*. *scapularis* and the parasitiform mites *D*. *gallinae*, *Tropilaelaps mercedesae* and *V. destructor* as initial queries in order to identify additional transcripts (Altschul et al., 1997). All of the candidate genes were manually verified by BLASTx against the non-redundant (NR) database at NCBI, and the open reading frames (ORFs) were identified using the ORFfinder at NCBI. We also tested signature patterns of Pfam domain architectures for IR/iGluRs (PF01094; PF00060; PF10613) and GRs (PF08395 and PF06151) using the HMMER software (Potter, et al., 2018). Thereafter, the amino acid sequences were blasted against the pfam (http://pfam.xfam.org/) database to obtain unigene annotation information (Finn et al., 2015). Lastly, prediction of membrane protein topology and signal peptides were performed with TOPCONS and TMHMM (Bernsel, et al. 2009). Amino acid sequences of IR/iGluRs predicted in both *O. sylviarum* and *D*. *gallinae*, along with *T*. *mercedesae*, *I*. *scapularis*, *Galendromus* (*=Typhlodromus, =Metaseiulus*) *occidentalis, V*. *destructor*, the vinegar fly *D*. *melanogaster*, and the crustacean *Daphnia magna* were aligned using MAFFT default settings. The GR sequences from *O. sylviarum* and *D*. *gallinae* were analyzed together with those from *T*. *mercedesae, I*. *scapularis*, *M*. *occidentalis, V*. *destructor*, *Tetranychus urticae*, *D*. *melanogaster*, and *Daphnia pulex*. The amino acid sequences used for phylogenetic tree construction are listed in Supplementary Material (Supplementary Table S1 and S2; Vizueta et al., 2018). A phylogenetic tree was performed with PhyML (maximum likelihood) software using the best-fitting likelihood model LG+G+I+F as calculated by SMS (Smart Model Selection) (Lefort et al., 2017; Katoh et al., 2019). A phylogenetic tree branch support was calculated using a new, fast, and approximate maximum likelihood-ratio test (aLRT), and the tree was visualized in EvolView (He et al. 2017).

### *Ab initio* modelling of protein sequence

The three-dimensional protein structure and function of selected proteins of *O. sylviarum* were predicted using an online version of I-TASSER (Iterative Threading Assembly Refinement) (Roy et al., 2010). The crystal structure of the AMPA subtype ionotropic glutamate receptor (PDB code: 3KG2) served as a template for the Osyl_iGluR7. For the Osyl_IR9, the crystal structure of *Rattus norvegicus* (PDB code: 5kuf) served as a template. Pymol was used to visualize the complex model. The quality of the resulting structural homology model was evaluated in comparison with the quality of the template.

## Results and discussion

### *De novo* assembly, functional annotation and classification of the *O. sylviarum* transcriptome

To gain relevant molecular information on *O. sylviarum*, as a mean to identify novel targets for pest control, we used the BGISEQ-500 next generation sequencing platform, which provided a total of 143.74 million raw reads (totaling roughly 19.72 GB of sequence data), of which 91.43 % were clean reads. Among the 131.41 million clean reads, 96.98% and 88.75 % had quality scores above the Q20 and Q30 levels, respectively (Supplementary Table S3). After Trinity *de novo* assembly of transcriptome, in total 28,999 unigenes were obtained with an average length of 1,635 bp. Total sequencing length, N50 and GC content were 47,424, 969 bp, 2,907 bp and 48.33%, respectively (Supplementary Table S4; Supplementary Figure S1). The extent of completeness and duplication of the *O. sylviarum* gene set were assessed using the arthropod-specific Benchmarking Universal Single-Copy Ortholog (BUSCO) genes (Simão et al., 2015). We retrieved 90.53% complete orthologs (C), being 24% duplicated (D) only 2.53% fragmented orthologs (F), and 6.94% missing (M) from the assembly (Supplementary Figure S2). BUSCO indicated a high degree of completeness at 93.06%, which suggests that most of our transcriptome assembly is significantly complete in terms of gene repertoire (Supplementary Figure S2). In total 19,750 unigenes were generated by the seven functional databases: 17,783 (61.32%) in NR, 6,353 (21.91%) in Nt, 12,956 (44.68%) in KOG, 13,528 (46.65%) in Swiss-Prot, 14,840 (51.17%) in Pfam, 5,853 (20.18%) in GO and 13,661 (47.11%) in KEGG annotation. The remaining 9,249 (31.89%) unigenes were not annotated, suggesting that some of them may represent lineage-specific novel genes, in addition to the typical presence of assembly artifacts or non-coding RNA. Similarly, 27,547 unigenes (67.86%) of the mite *D*. *gallinae* were classified as unknown function, highlighting the need for functional genomic studies for poultry mite species (Huang et al., 2020).

Based on the unigenes with GO annotation (20.18%), these unigenes were primarily divided into three separate ontologies: biological processes, molecular functions and cellular components. The most abundant categories were classified into catalytic activity (2,667 unigenes), binding (2,620 unigenes) and cellular processes (1,950 unigenes) (Supplementary Figure S3). Apart from the GO ontology analysis, a total of 13,661 (47.11%) unigenes were used for KEGG analysis, which identified six main pathway branches (Supplementary Table S5). Among these, ‘metabolism’, ‘environmental information processing’ and ‘cellular process’ were the most dominant categories, in which metabolism related pathways occupied the bulk part in NFM. These results can provide a valuable reference for investigating specific biological, molecular functions and pathway analyses of *O*. *sylviarum* transcriptome (Supplementary Figure S4). According to the NR database, the top-hit species was the honey bee parasitic mite *T*. *mercedesae* (63.88%), followed by the predatory mite *G*. *occidentalis* (24.72%) and the deer tick *I*. *scapularis* (0.59%), suggesting that the NFM transcriptome matches well with other Parasitiformes mites. Thus, poultry mites share most of the orthologous genes with the parasitiform mites *T. mercedesae* and *G. occidentalis* rather than with the parasitiform tick *I*. *scapularis* (Figure 1), which is in agreement with a previous study (Huang et al., 2020). The KOG analyses can provide useful information of the annotated unigenes based on gene-based definition of orthology. In total of 44.68% unigenes with non-redundant database hits were grouped into 25 KOG categories (Supplementary Figure S5). For *O*. *sylviarum*, the largest categories were ‘general function prediction only’, followed by ‘signal transduction mechanisms’, ‘function unknown’, and ‘posttranslational modification, protein turnover’. The smallest unit of classification were ‘cell motility’, “nuclear structure’ and ‘nucleotide transport and metabolism’. The raw reads have been deposited in the Sequence Read Archive (SRA) database under the accession number PRJNA675700, while the Transcriptome Shotgun Assembly (TSA) has been submitted with accession number GIXZ00000000.

**Figure 1:**
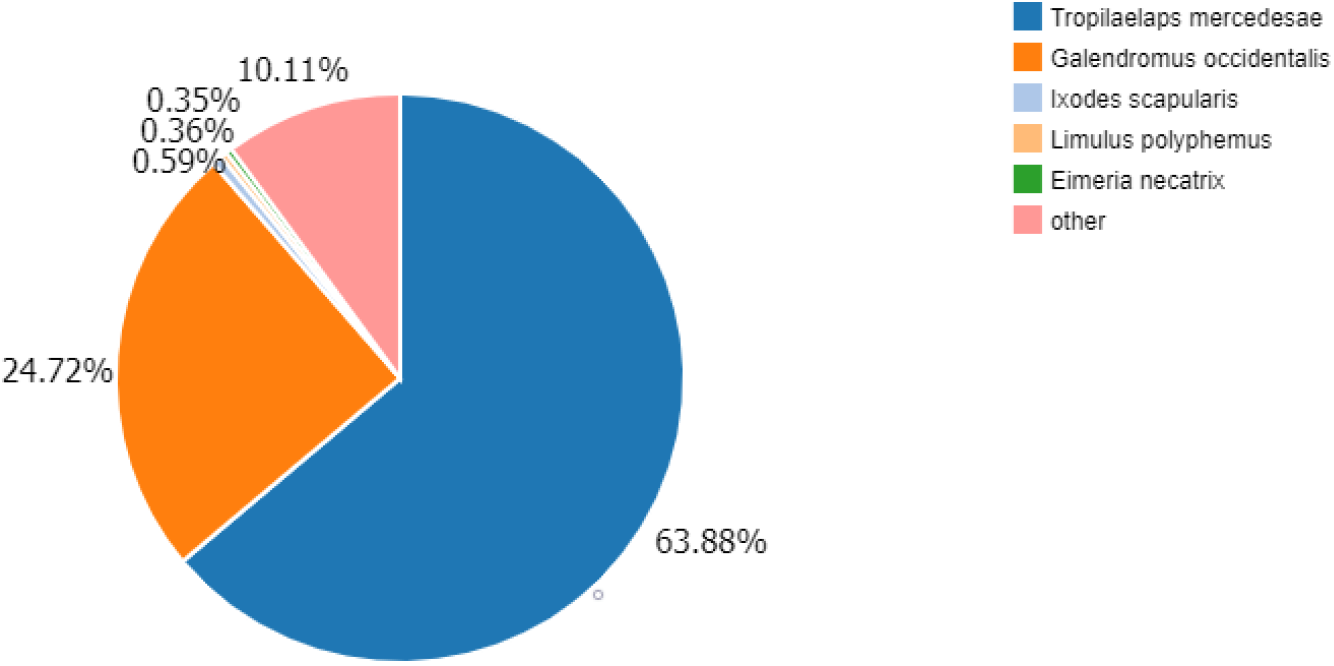
The species distribution of BLASTx matches of the transcriptome unigenes. Each section shows the number of top BLASTx hits.

### Loss or reduction of highly conserved transcription factors in *O. sylviarum*

Transcription factors (TFs) are key proteins that bind specifically to DNA, and are an integral part of gene expression at the transcriptional level. Currently, there is little information available for the structures and biological functions of TFs in mites. In the present study, 2,191 putative unigenes were identified and found to be distributed across 65 TF families. Among these, zf-C2H2 was the most abundant family (668 unigenes), followed by Homeobox (172 unigenes), and bHLH (125 unigenes) (Figure 2). Homeobox genes were present in a range of arachnids that contain a highly conserved homeodomain. Widespread retention and pervasive gene duplication of homeobox gene has been observed in Arachnida class (e.g., spiders, ticks, scorpions, and mites), suggesting implications for the field of evolutionary biology (Leite et al., 2018). Genomic analyses have also reported that mite species exhibit dynamic rearrangements of homeobox clusters, along with loss of highly conserved TFs (Hoy, 2009; Leite et al., 2018; Grbic et al., 2011). The basic helix-loop-helix (bHLH) is one of the largest family of transcriptional regulatory proteins, and are present in organisms from yeast to humans, where their members are participating in regulating a wide range of biological and developmental processes, such as the development of the nervous system and muscles, as well as responding to environmental factors (Jones, 2004). Notably, a number of specific TFs that are highly conserved in most arthropods were not detected in the *O*. *sylviarum* transcriptome, including circadian rhythm (so-called Clock-Cycle), E78, HR3, Har-AP-4, HR39, EcR, Abd-A, ERR, Ato, FTZ-F1, ATF-3, HR96, Dip3, NRs HNF4, HR78, CREB, HR83, and Dnato3 (Guo et al., 2018). The absence of these TFs is highly intriguing, and clearly demands further investigation, using different transcriptome or genomic data in order to validate this result.

**Figure 2:**
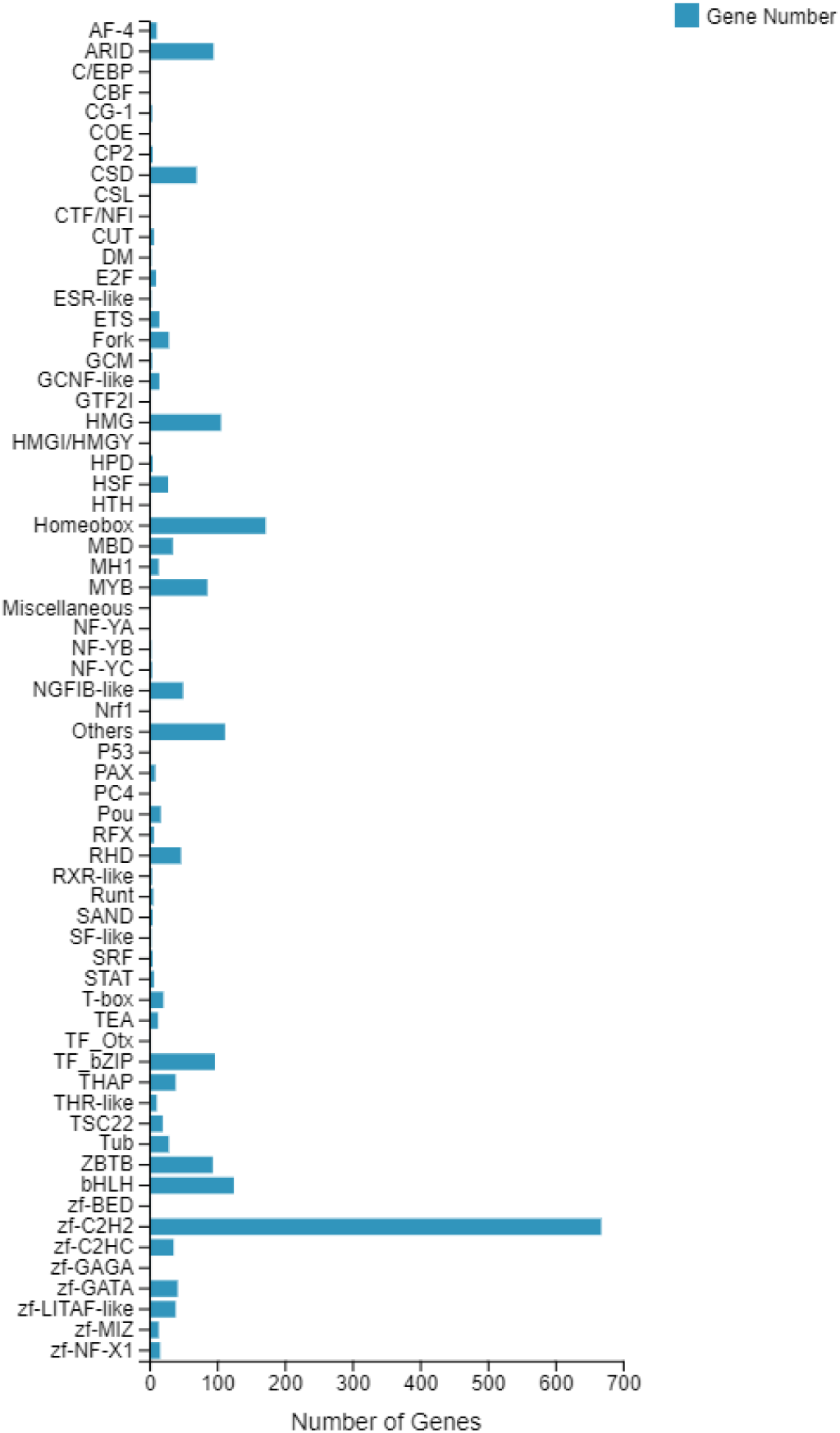
Family distribution of the transcription factors (TFs) in the NFM transcriptome. The number of genes identified in each TF family are represented in X-axis.

### Identification and interactive web-based annotation of chemosensory receptors

Annotation of the IRs and GRs by a bioinformatics-based approach, coupled with extensive manual curation, revealed only 1 and 2 receptors in the NFM transcriptome, respectively. The major limitation of the study is that, the transcriptome data of NFM is generated from a pooled sample comprising all life-stages and sexes, and thus their transcripts are likely to be highly diluted in the pooled sample. It is also important to note that the number of candidate chemosensory receptors identified in NFM is similar to that found in a recent transcriptome analysis of chemosensory gene expression in different organs in PRM (Bhowmick et al., 2020), but considerably smaller than that described in other free-living mite and tick species, as well as in insects (Figure 3). Relying on only transcriptomic data, however, may not be suitable to assess the presence or absence of IRs and GRs. Some of these receptors including GPCRs (G-protein coupled receptor) are difficult to identify in the transcriptome assembly because they are present in specialized sensory tissues (e.g. Haller’s organ in ticks and foretarsal sensory organ in mites) and lower level of gene expression profiles (Bhowmick et al., 2020; Pietrantonio et al., 2018; Vizueta et al., 2020b).

**Figure 3:**
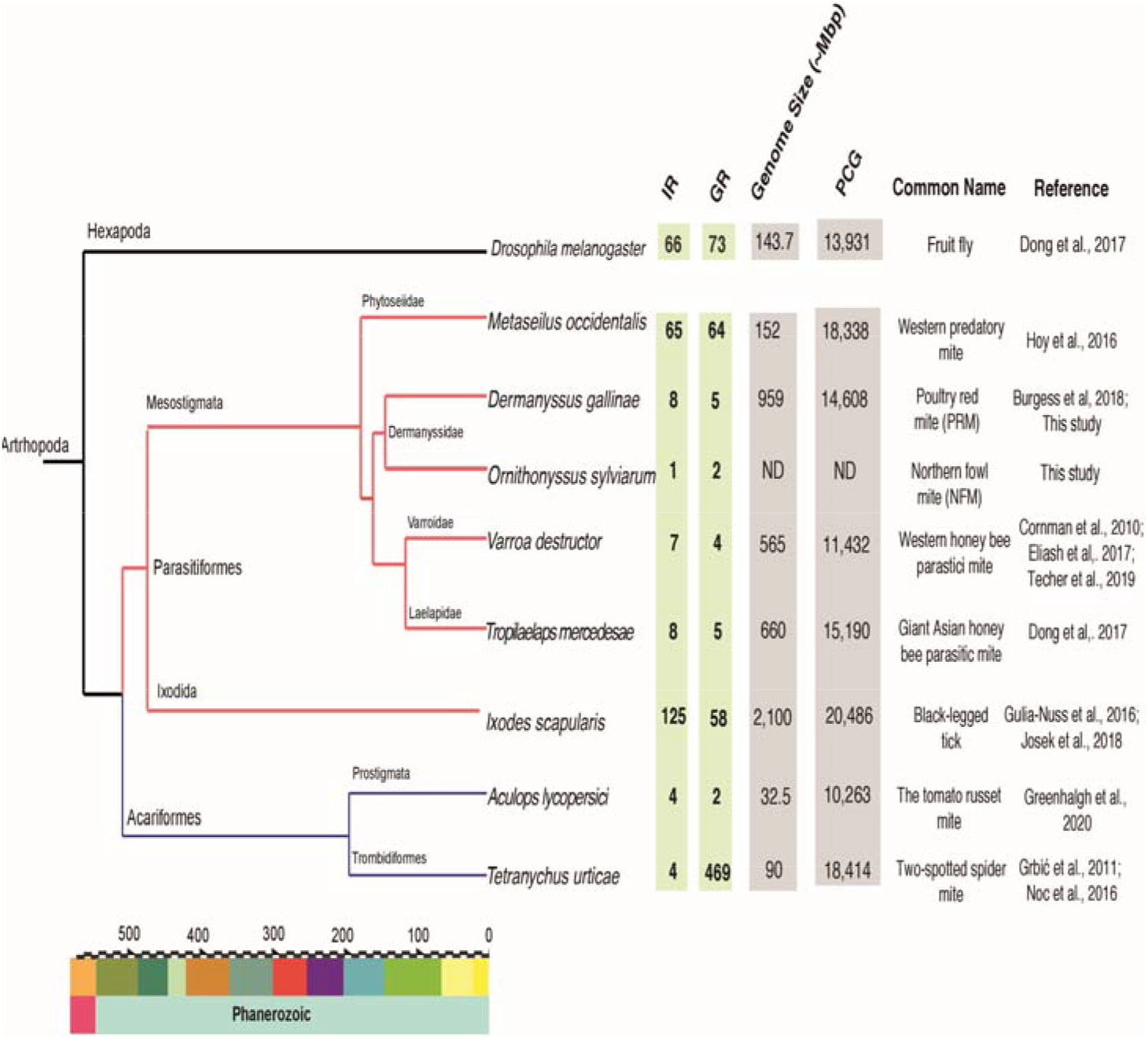
The number of candidate chemosensory receptor genes, genome and transcriptome statistics of poultry mites and other arthropods species. To compare the differences of IR and GR gene numbers among various mite and tick species, we compared two species of poultry mite chemoreceptor genes to those of three other Mesostigmata species and those of three additional Acari and an insect. Species are depicted with evolutionary relationships and divergence times based on Lozano-Fernandez et al. (2020) and Howard et al. (2020), with the timescale axis representing millions of years before the present. PCG: protein coding genes; ND: not determined; Mbp: million base pair.

To assess if the specialized lifestyle (extreme host specificity) of NFM has had a notable impact on the presence and abundance of IRs and GRs, we included the complete genome of the PRM, which has a similar lifestyle, in further analysis, and compared this dataset with the well-annotated genome of *I*. *scapularis*, which exhibits a different lifestyle (a broad range of hosts, including humans) (Figure 3). Nine GRs and forty-one iGluRs/IRs genes were annotated in the PRM genome, which have increased the completely new chemosensory gene regions in the NCBI database for this species. Accurate gene annotation from newly sequenced genomes is an important step for downstream functional and evolutionary analyses. Here, we used the BITACORA package to identify and predict all chemoreceptor family members, and then validated the chemosensory genes via ORCAE. The ORCAE-lead platform provides full annotation information about the gene models, such as gene structurs, location of genes in the genome, visualization of gene expression atlas, protein homologs and domains, and definition of gene function. Each gene ID was assigned a unique loci identifier with the following format: DEGALXgY, where DEGAL denotes the species name (*D*. *gallinae*), X indicates the scaffold ID and Y defines the specific location within the scaffold. The predicted gene models were generated using both a computational stand-alone pipeline and manual curation, along with global gene expression profile or EST alignment data from ORCAE, which means that almost all olfactory gene loci were identified at correct locations (Figure 4). As such, annotation of the *D*. *gallinae* draft genome can be further improved via a process of community-led manual curation.

**Figure 4:**
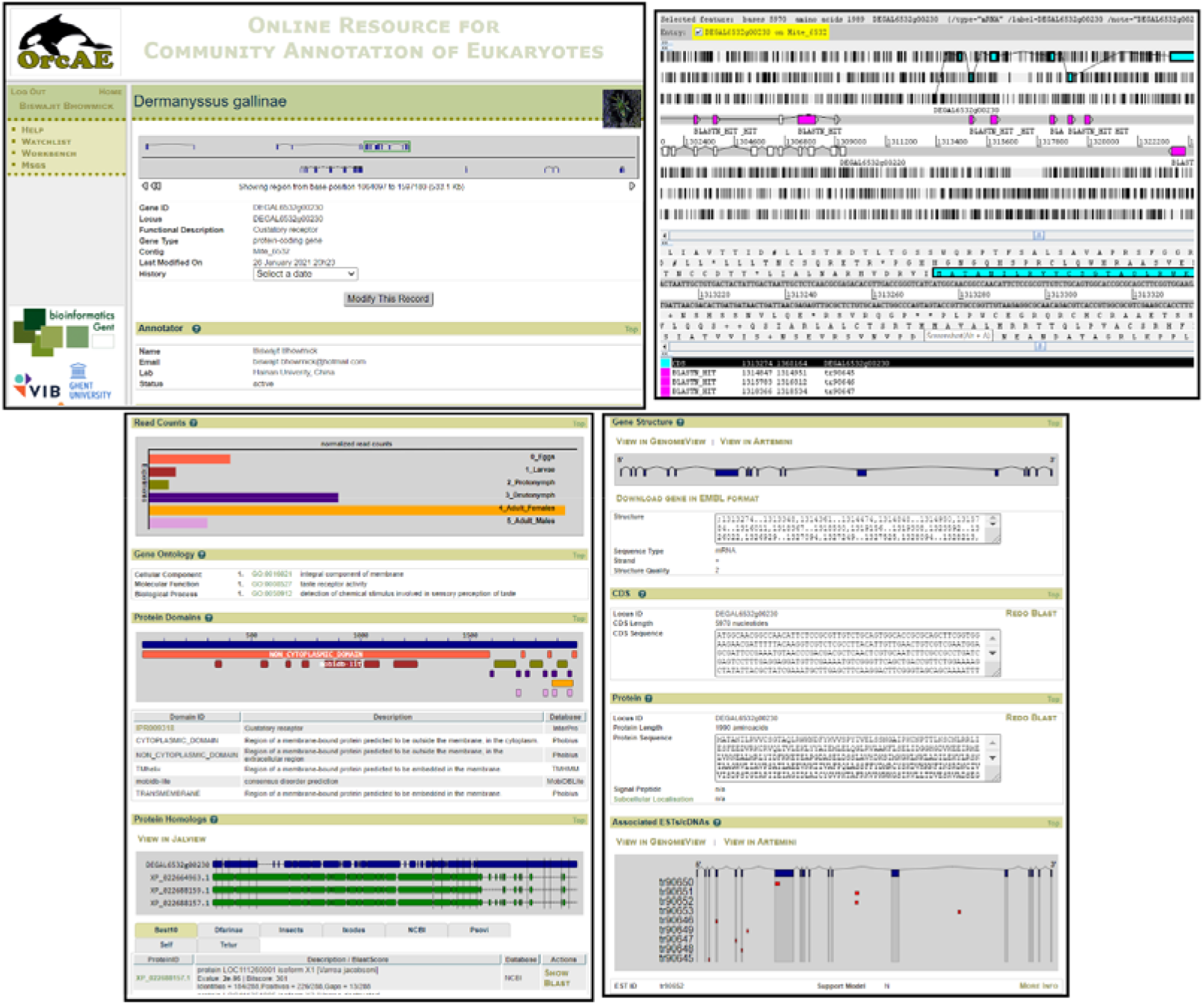
An overview of gene pages in the ORCAE resource. These pages display an extensive graphical representation of gene structure, function, coding sequence (CDS), orthologous information from other public databases, protein domains for a given gene, as well as other evidential information such as expressed sequence tag (EST) and gene expression profiles. The gene-specific page for the gustatory receptor (GR), DEGAL6532g00230 is displayed. DEGAL means the species name (*Dermanyssus gallinae*), 6532 denotes the scaffold ID and 00230 defines the specific location within the scaffold. Expression of this GR gene was supported by EST data.

### Sequence and phylogenetic analyses of candidate IRs/iGluRs and GRs

Out of nine candidate genes encoding IR/iGLuRs genes in *O*. *sylviarum*, six unigenes contained the specific domain signature of the ligand-gated ion channel (LCD-PF00060), which is characteristic of most IR/iGluRs subfamilies (Croset et al., 2010). Only two candidate unigenes had an amino terminal domain (ATD-PF01094), which is characteristic of traditional iGluRs (AMPA, NMDA and kainate receptors) (Croset et al., 2010). In *D*. *gallinae*, a total of 41 IRs/iGLuRs unigenes were predicted in the genome assembly, of which 29 unigenes contained LCD domain. To obtain additional evidence supporting the homology of the conserved family of iGluR, we carried out *ab initio* 3D structure predictions of Osyl_iGluR7 (AMPA-type). Osyl_iGluR7 has a unique modular architecture with four distinct domains: (1) an ATD, which is structurally and functionally the most divergent site among the iGluR subunits, is involved in receptor assembly, trafficking and modulation; (2) a ligand-binding domain (LBD), which is involved in agonist and/or antagonist recognition to activate ion channel; (3) a transmembrane domain (TMD), which forms the membrane-spanning ion channel; and (4) an intracellular C-terminal domain (CTD) involved in regulating synaptic efficiency, receptor mobility, and trafficking (Figure 5). In order to classify the IR/iGLuR family *O*. *sylviarum* and *D*. *gallinae*, we inferred a maximum-likelihood phylogenetic tree based on domain-specific region analysis (PF00060), which identified four major clades. Of the seven candidate IR/iGluRs unigenes, with a conserved domain, four (Osyl_iGluR1, Osyl_iGluR2, Osyl_iGluR3, Osyl_iGluR4) were members of the kainate receptor subfamily, one (Osyl_iGluR7) was phylogenetically clustered with AMPA receptors, and one (Osyl_iGluR8) clustered in an NMDA receptor subfamily (Figure 6). A similar number of candidate iGluR unigenes were identified in the genome of *D*. *gallinae*: 10 kainate-type, 5 AMPA-type and 2 NMDA-type receptors (Figure 6). Our phylogenetic analysis also revealed more divergent Kainate-type orthologs among three classical iGluRs. This expansion is possibly derived from gene duplications.

**Figure 5:**
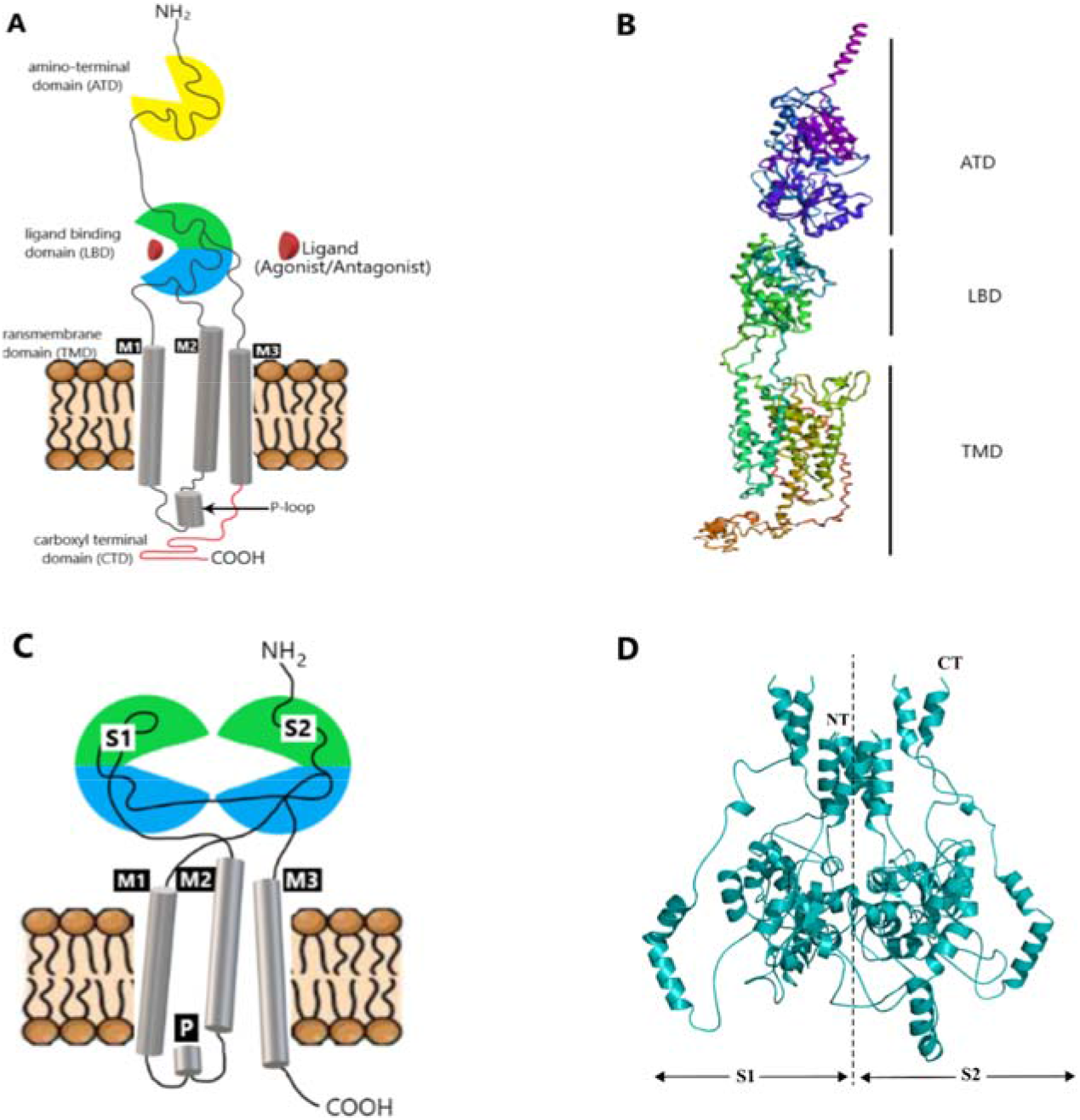
Overall topology of the ionotropic glutamate receptor (iGluR) and ionotropic receptor (IR). (A) A schematic representation of the iGluR protein is shown in cartoon form. The two halves of LBD (S1 and S2) are present; M1, P loop, M2, and M3 make up the transmembrane (7_tm) region of each subunit. (B) The homology model for the 3D structure of Osyl_IR7 (AMPA-type iGluR) was created via I-Tasser. Predicted structure of the putative AMPA-type receptor demonstrates distinct domains, including the amino terminal domain (ATD), the ligand-binding domain (LBD) and the transmembrane domain (TMD). The intracellular carboxyl-terminal domain (CTD) is not shown. (C) A general schematic representation of IR protein is demonstrated in cartoon form. ATD is absent in most IRs. The two halves of LBD (S1 and S2) are present. (D) The homology model for the 3D structure of Osyl_IR9 (IR25a-like) is shown. The bilobal domain architecture is marked by the dashed line, with the S1 domain in the left and the S2 domain in the right side. NT means N-terminal region, and CT indicates C-terminal region.

**Figure 6:**
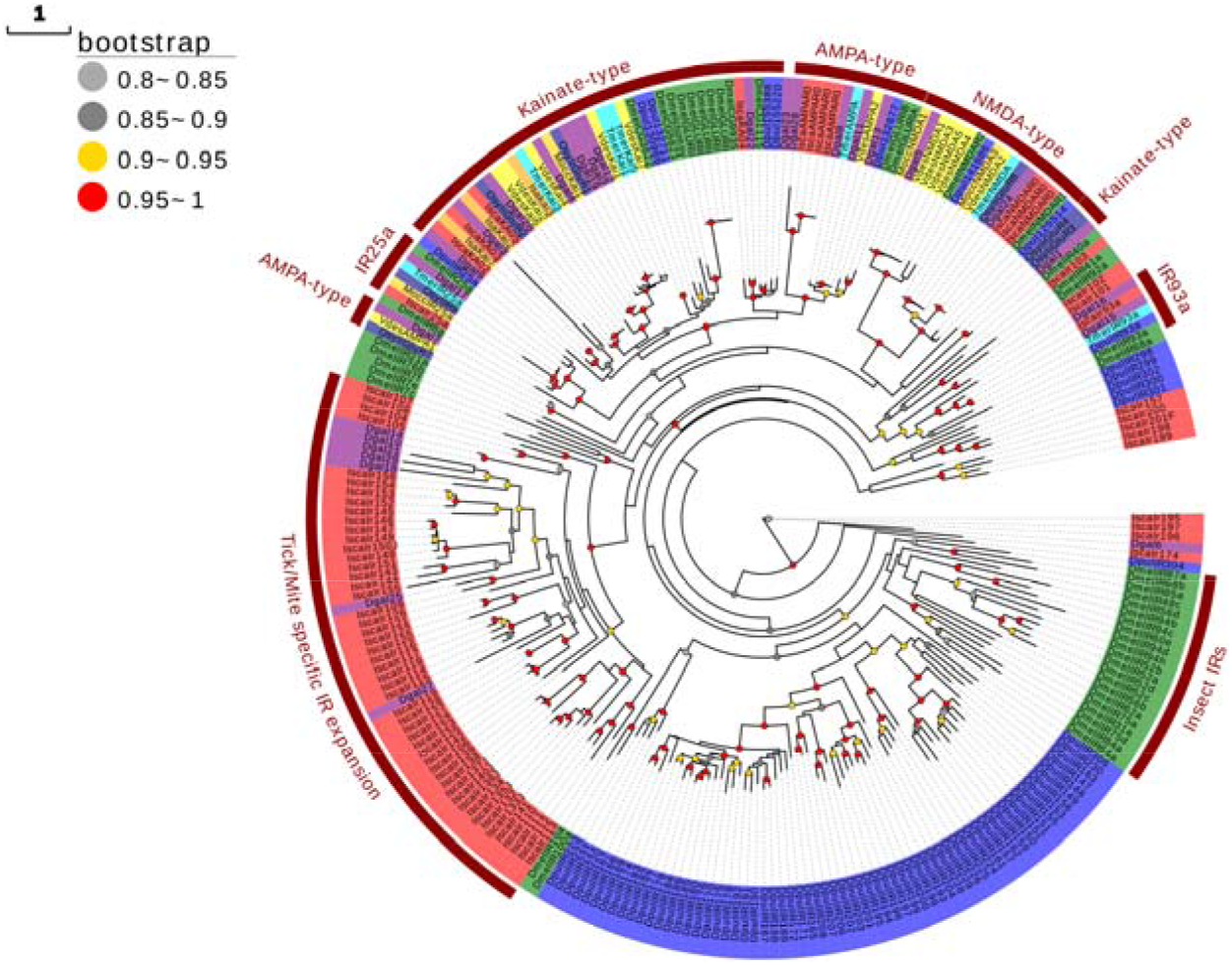
An evolutionary tree of the iGluR/IR gene family is based on LCD domain. This analysis involved 265 protein sequences from *Ornithonyssus sylviarum* (dark blue: Osyl), *Dermanyssus gallinae* (purple: Dgal), *Varroa destructor* (yellow: Vd), *Metaseiulus occidentalis* (gold: Mocc), *Ixodes scapularis* (red: Isca), *Tropilaelaps mercedesae* (cyan: Tmer), *Drosophila melanogaster* (green: Dmel) and *Daphnia pulex* (blue: Dpul). The sequences that contain the relevant conserved domain (PF00060) were retrieved and used to construct phylogenetic analysis. The scale bar indicates amino acid changes per site.

Like iGluRs, IRs are characterized by a presence of short extracellular N-terminal region, a bipartite LBD (S1 and S2), LCD and a short C-terminal region. However, our *ab initio* protein structure prediction from the amino acid sequence (Osyl_IR9; IR25a-like) did not identify LCD because the homologous proteins that have not been solved experimentally (Figure 5). In spite of having a similar domain architecture to iGluRs, amino acid sequence similarities between IRs and iGluRs is quite low (less than 34%), particularly within the LBD. Most IRs that lack an ATD require a co-receptor to function (Rytz et al., 2013; van Giesen et al., 2017). Apart from the iGluRs, the conserved co-receptor IR25a was identified in both *O. sylviarum* and *D*. *gallinae*, which is the oldest member of the IRs (Croset et al., 2010) (Figure 6). Interestingly, IR25a from both species shows high sequence similarity with IR25a identified in distantly related species, e.g. the vinegar fly *D. melanogaster*. Orthologs of the other conserved IR co-receptors for olfactory function, IR8a and IR76b (Croset et al., 2010; Vizueta et al., 2018), that are present in the genomes of most insects and non-insect arthropods, were not identified in *O. sylviarum*, the phylogenetic analyses confirmed the presence of two putative IR93a members in *D*. *gallinae*, as well as 8 complete divergent IRs (Figure 6). While the transcriptome of *O*. *sylviarum* is highly complete according to the BUSCO searches, the absence of co-receptors, other than the IR25a ortholog, and divergent IRs in in this species may be caused by the fragmentation of sequences in the assembly, or by the lack of expression of these gene family members in our transcriptome. The presence of a few orthologous IR co-receptors and divergent IRs in other mites (Figure 3), along with our assessed species, suggests either that the last common ancestor of arthropods had very few IRs, or that IRs may have been lost due to different adaptations. In support of this, Parasitiformes mites that are exposed to different types of environments and multiple host range, such as in the case of the predatory phytoseiid mite and the black-legged tick, have a higher number of IRs than the more strictly host-specific poultry mites (Figure 3).

Two candidate GRs, with complete ORFs containing the seven transmembrane domains (PF08395), were identified in the *O*. *sylviarum* transcriptome, whereas a total of nine GR genes, presenting either the 7m_7 (PF08395) or trehalose (PF06151) domains (Zhang et al., 2011), were found in the PRM genome. These candidate genes were used in phylogenetic analysis, together with representatives of the three most reported GR gene family, i.e., those involved in the detection of CO_2_, bitters and sugars (Zhang et al., 2011). While we were unable to identify any orthologues of these GR within these lineages in poultry mites, this may not be surprising as these genes are barely identifiable in nono-holometabolous insect lineages, including the dampwood termites *Zootermopsis nevadensis* and the damselflies *Calopteryx splendens* (Terrapon et al., 2014; Ioannidis et al., 2017) (Figure 7). Most of the mite GRs clustered with GRs of ticks, suggesting that the GR repertory in these organisms originated from gene expansions specific to Acari. This suggests that Acari might use phylogenetically divergent GRs for taste detection. Chelicerates (e.g., spiders, ticks, scorpions, and mites) have evolved a lineage-specific set of GR expansions not shared with other arthropods, as expected from their fast evolutionary rates and rapid sequence evolution (Vizueta et al., 2018). Interestingly, among the nine GRs in the *D*. g*allinae*, one GR (DEGAL5202g00060.0) was grouped with bitter receptor from *D*. *melanogaster* with strong bootstrap support (◻ 0.95). Future functional studies are needed in order to shed light on the specific functions and evolution of chemoreceptors in Acari.

**Figure 7:**
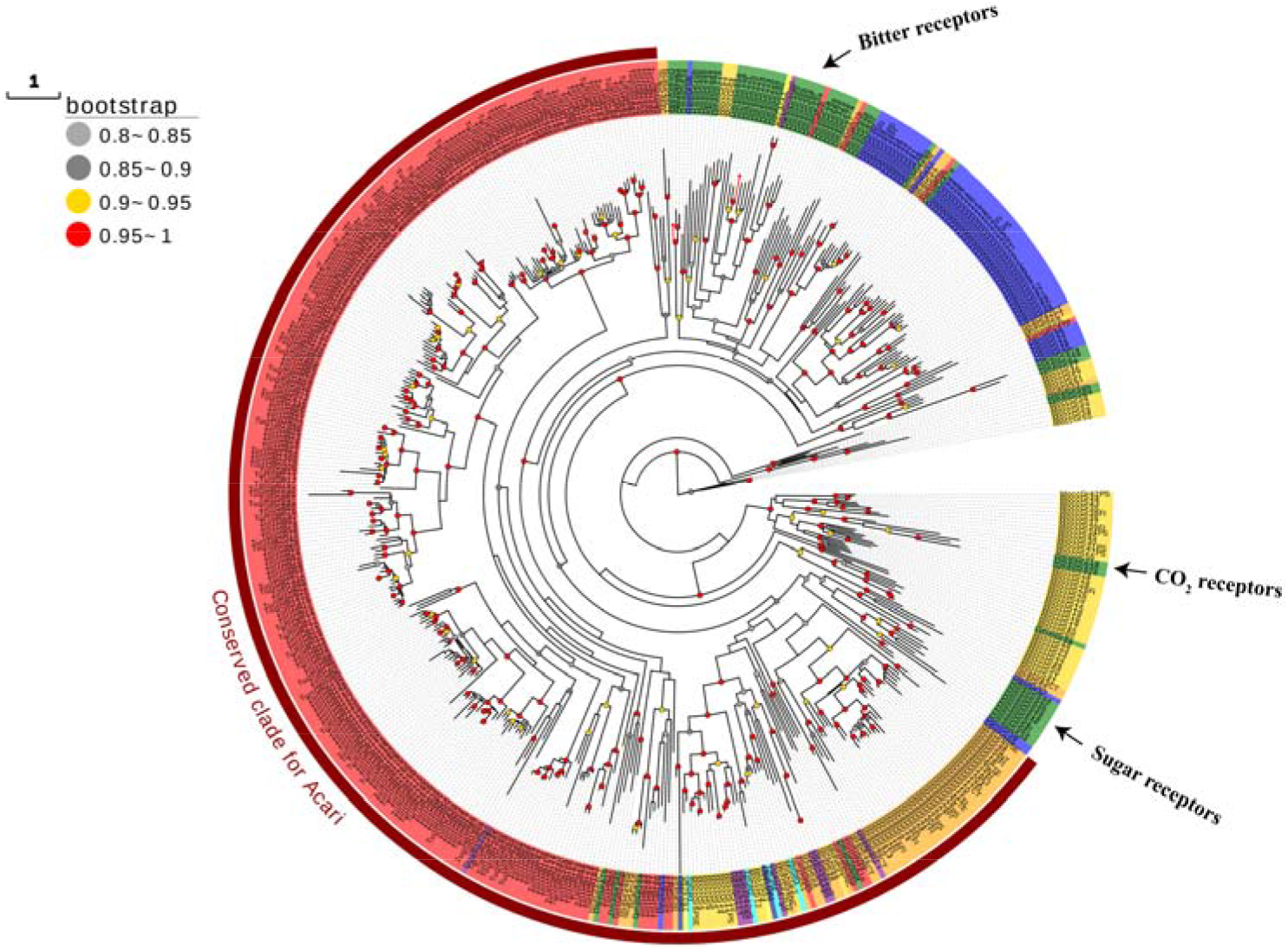
A phylogenetic tree of the GR proteins across arthropods is shown. This analysis involved 518 GR protein sequences from *Ornithonyssus sylviarum* (dark blue: Osyl), *Dermanyssus gallinae* (purple: Dgal), *Varroa destructor* (yellow: Vd), *Tetranychus urticae* (red: Turt), *Metaseiulus occidentalis* (gold: Mocc), *Ixodes scapularis* (orange: Isca), *Tropilaelaps mercedesae* (cyan: Tp), *Drosophila melanogaster* (green: Dmel) and *Daphnia pulex* (blue: Dpul). The scale bar indicates amino acid changes per site.

### Fewer chemosensory genes are associated with the life history of poultry mites

Both *O. sylviarum* and *D*. *gallinae* have significantly lower numbers of predicted IR and GR genes in comparison with *I*. *scapularis* (Figure 3). This is likely to be explained by their simple chemosensory system of poultry mites either due to their “lifestyle as obligate parasites”, or the “result of a lack of duplication”. The former describes the evolutionary force that possibly shaped the genomic changes, whereas the latter refers to a molecular mechanism through which the changes may occur. This is in line with that described for the the human body louse, *Pediculus humanus humanus,* the bed bug, *Cimex lectularius* and the tomato russet mite, *Aculops lycopersici* (TRM; the world’s smallest plant-eating mites), which all have a low number of genes associated with chemoreception as well as detoxification (Kirkness et al., 2010; Benoit et al., 2016; Greenhalgh et al., 2020). Alternatively, the small chemosensory gene set may be a result of a lack of duplication (fewer expansions of presumably small ancestral gene families repertoires), which previously has been demonstrated in *T*. *mercedesae*, which has undergone the fewest gene family expansion or contraction events since its divergence from the last common ancestor of arthropods (Dong et al., 2017). A lower number of genes associated with environmental sensing have also been shown to be present in other obligate parasites, compared to closely related free-living relatives in flies, ticks and beetles (Kirkness et al., 2010). As a whole, comparative genomics and transcriptomics analysis offer unique information and tools to use in advancing the understanding of host-parasite co-evolution and signatures of adaptive evolution. Further functional studies combined with new evidence based on greater coverage of phylogenetic analyses can greatly contribute to understand the chemosensory gene evolution.

## Conclusions

In conclusion, the *O*. *sylviarum* transcriptomic data and annotation results can provide a useful resource for future studies in this non-model mite species, which is an economically important pest. This study improves our understanding of its molecular biology that may result in an improved integrated pest management program, providing a tool for further identification and characterization of genes of interest and development of eco-friendly acaricides or mite repellents. Additionally, a comprehensive analysis of variant and conserved receptors in two species of poultry mites revealed many features that will help in elucidating the molecular basis of the function and regulation of the chemosensory system in mites. The decreased number of chemosensory receptors predicted in bird mites may reflect the unique adaptations of these species to their specific behavioral and reproductive lifestyle.

## Supplementary material

The Supplementary Material for this article is provided.

## Ethics statement

Not applicable.

## Author contributions

BB and QH conceived and designed the study. BB and HC wrote the first draft and performed the experiments. JL, JV, RI, and QH commented on study design, methodology, and substantially revised the manuscript. All authors approved the final version.

## Acknowledgments

We would like to thank Yu Tang, Dr. Jianguo Zhao, and Dr. Chenghong Liao for their assistance. This work was supported by the National Key Research and Development Program of China (2017YFD0501702-9) and Hainan Provincial Key R & D Program (ZDYF2019073).

